# ALPPL2 is a highly specific and targetable tumor cell surface antigen

**DOI:** 10.1101/2020.01.07.898122

**Authors:** Yang Su, Xin Zhang, Scott Bidlingmaier, Christopher R. Behrens, Nam-Kyung Lee, Bin Liu

## Abstract

It has been challenging to identify tumor-specific cell surface antigens as the vast majority of tumor-associated antigens are also expressed by some normal tissues. In the course of our study on mesothelioma, we identified a highly specific tumor cell surface antigen that can be targeted for therapy development. Mesothelioma is caused by malignant transformation of the mesothelium, incurable and categorized into three histological subtypes, epithelioid, biphasic and sarcomatoid. To identity novel mesothelioma cell surface antigens with broad subtype coverage and high tissue specificity, we have previously selected phage antibody display libraries on live mesothelioma cells and tissues following counter-selection on normal cells, and identified a panel of human antibodies that bind all subtypes of mesothelioma but not normal mesothelium. One of the antibodies, M25, showed high specificity, and we hereby report the identification of the M25 antigen as ALPPL2. We performed immunohistochemistry on normal human tissues and found that ALPPL2 is expressed only on placental trophoblasts but not any other normal tissues. This exquisite tissue specificity and broad tumor type coverage suggests that ALPPL2 could be an excellent cell surface target for therapeutic development against mesothelioma. To evaluate therapeutic potential of ALPPL2 targeting, we developed an ALPPL2-targeted antibody-drug conjugate and demonstrated potent and specific tumor killing *in vitro* and *in vivo* against both epithelioid and sarcomatoid mesothelioma. Thus ALPPL2 belongs to a rare class of cell surface antigens that can be said as being truly tumor specific and is well suited for therapy development against ALPPL2 expressing tumors.

## Introduction

Mesothelioma is a rare, asbestos-related tumor (Wagner et al., 1960) that arises predominantly from serosal cells of the pleura and, to a lesser extent, the peritoneum, pericardium, and tunica vaginalis (Yap et al., 2017). About 3,000 new cases are diagnosed annually in the United States (Enewold et al., 2017; Zervos et al., 2008). Current approved treatments for malignant mesothelioma include chemotherapy, radiation therapy, or combination treatments (Katzman and Sterman, 2018; Zervos et al., 2008). However, malignant mesothelioma is incurable, and the overall five-year survival rate is about 8% (Abdel-Rahman et al., 2018; Taioli et al., 2017).

Malignant mesothelioma can be classified into three histological types, epithelioid (∼50-60%), sarcomatoid (fibrous, ∼10-20%), and mixed (biphasic, ∼30-40%) (Galateau-Salle et al., 2016; Yap et al., 2017). Epithelioid mesothelioma tends to have a better prognosis than the other two types. Sarcomatoid mesothelioma is particularly recalcitrant to treatment, progresses most aggressively and has the shortest median overall survival (Muruganandan et al., 2017; Yap et al., 2017). Currently, the best studied cell surface antigen for mesothelioma is mesothelin, a GPI-anchored molecule, which is the only biomarker approved by the FDA for this disease (Hassan et al., 2016; Li et al., 2007). Mesothelin, however, is rarely expressed by sarcomatoid mesothelioma (Ordonez, 2003). In addition, mesothelin is also expressed by normal mesothelium, raising concerns for on-target toxicity on normal tissues. Therefore new targets are needed for developing novel therapies against this incurable malignancy.

An ideal cell surface target for mesothelioma should bear the following desired properties: (1) it is expressed in all subtypes of mesothelioma but not normal tissues. (2) it is a true tumor antigen that is expressed by mesothelioma cells *in situ* residing in their tissue microenvironment, as opposed to cell line artifacts. We have previously developed methods for selecting phage human antibody display libraries directly on tumor issues derived from patient specimens (An et al., 2008; Bidlingmaier et al., 2009; Ruan et al., 2006) following counter-selection on normal tissues and cells. In the case of mesothelioma, we identified a panel of novel human a single-chain variable fragments (scFvs) that bind to both epithelial and sarcomatous types but not normal mesothelium (An et al., 2008), a clear differentiation from antibodies binding to mesothelin that is expressed by epithelial mesothelioma but rarely by sarcomatous mesothelioma (Ordonez, 2003), and also by normal mesothelium (Petricevic et al., 2012). One of the antibodies, M25, showed the highest specificity, binding to a novel cell surface target expressed by all subtypes of mesothelioma (An et al., 2008).

We hereby report the identification of the target antigen bound by M25 as human alkaline phosphatase, placental-like 2 (ALPPL2), a member of the human alkaline phosphatase family. Among the four members of this family, ALPPL2 and placental alkaline phosphate (ALPP) are virtually identical in amino acid sequence (98% homology) and have a highly restricted tissue expression pattern, expressing in placental trophoblasts only. Both share high homology with the intestinal alkaline phosphatase (ALPI) (87% homology), and some homology with the tissue-nonspecific liver/bone/kidney phosphatase ALPL (57% homology) (Henthorn et al., 1988). We show that M25 binds specifically to ALPPL2 and ALPP but not ALPI or ALPL. We performed extensive immunohistochemistry (IHC) studies and showed that ALPPL2 is expressed in mesothelioma but not any other normal tissue except for placental trophoblasts, thus demonstrating an exquisite tissue specificity. ALPPL2 is therefore one of those rare cell surface antigens that can be classified as being truly tumor specific. To evaluate ALPPL2 as a potential therapeutic target, we constructed antibody-drug conjugates (ADCs) by conjugating microtubule inhibitors to our anti-ALPPL2 human monoclonal antibody M25 and showed that M25 ADCs potently inhibited tumor cell proliferation *in vitro* and mesothelioma cell line xenograft growth *in vivo*. We propose that ALPPL2 is a novel tumor cell surface target and its high tissue specificity makes it an excellent target for various forms of anti-cancer therapy.

## Results

### Identification of the antigen bound by M25 as ALPPL2

We have previously identified by phage display a novel scFv M25 that binds to both epithelioid and sarcomatoid subtypes of mesothelioma, but not the normal mesothelium (outlined in Figure 1A, ref. (An et al., 2008)). To identify the cell surface antigen bound by M25, we labeled cell surface proteins by biotinylation of live mesothelioma cell surface, and performed immunoprecipitation (IP) using matrix-immobilized M25 scFv-Fc followed by mass spectrometry analysis (outlined in Figure 1A, ref. (Ha et al., 2014)). As shown in Figure 1B, one band running at about 72 kDa on SDS-PAGE was identified from IP products of the mesothelioma cell line M28, but not the control benign prostatic hyperplasia (BPH-1) cell line. The corresponding band on duplicated SDS-PAGE gel was excised, in-gel digested, analyzed by tandem mass spectrometry, and identified as human ALPPL2. The theoretical molecular weight of ALPPL2 is about 57 kDa. Due to post-translational modifications, the apparent molecular weight of ALPPL2 is about 70 kDa (Watanabe et al., 1992).

**Figure 1.**
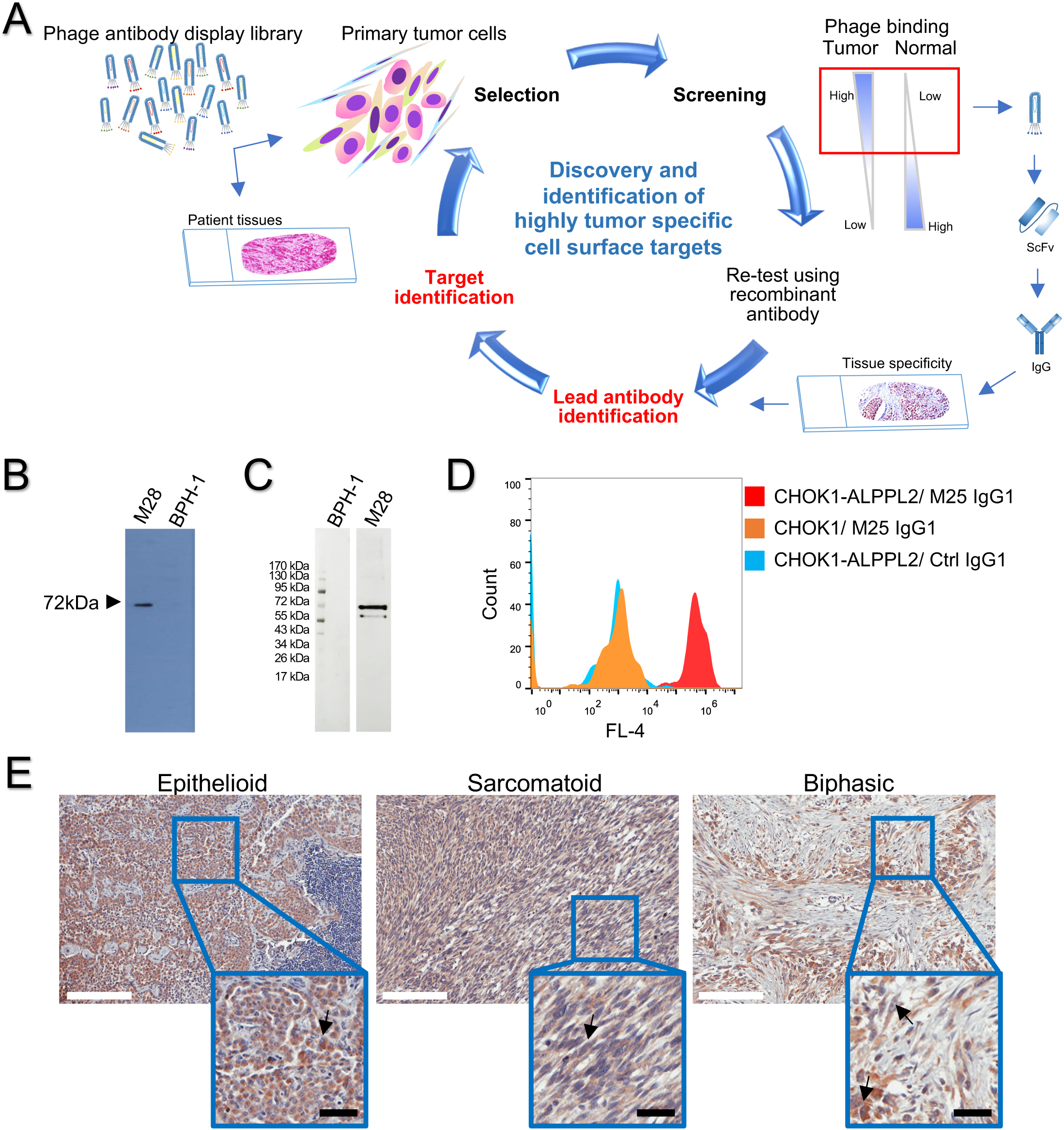
Identification of ALPPL2 as a highly tumor specific cell surface antigen. (A) Outline of approach that was used to discover and characterize tumor specific cell surface targets using phage antibody display library selection. Naïve phage display antibody library was incubated with primary tumor cells or tumor tissue slides following counterselection on normal cell/tissue. Tumor selective binders were screened for differential binding to a panel of tumor and normal control cells. Variable regions of tumor-specific phage binders were cloned to generate scFvs and IgGs for various characterizations including tissue specificity by IHC. Once a lead antibody is identified, it is used for target identification by immunoprecipitation from surface-biotinylated tumor cell lysates followed by mass spectrometry analysis. (B) Identification of the target antigen bound by the tumor specific antibody M25. Beads coated with M25 scFv-Fc were used to capture the target antigen from surface biotin-labeled M28 cell lysates, and the IP product was analyzed by Western blot using streptavidin-HRP. One band at 72 kDa is seen from IP products from tumor (M28) but not control nontumorigenic cells (BPH-1). (C) Verification of ALPPL2 as the target. M25 IP products from M28 and BPH-1 cells were probed with a commercially validated anti-human ALPPL2 antibody by Western blot. The anti-ALPPL2 antibody recognizes a 72 kDa antigen in the IP product from M28 but not BPH-1 cells. (D) M25 IgG1 binds to CHO-K1 cells ectopically expressing ALPPL2. CHO-K1 cells were transiently transfected with ALPPL2 expression plasmid, and binding of M25 IgG1 was analyzed by flow cytometry. (E) IHC study of FFPE mesothelioma tissue samples. Positive ALPPL2 staining is seen for different subtypes. Scale bar (white): 200 μm. Enlarged windows are close-up views. Arrow points to a representative tumor cell (not all tumor cells are marked). Scale bar (black): 30 μm.

To further validate the finding, we repeated the M25 IP experiment using M25 scFv-Fc-conjugated beads and probed the IP product with a commercial anti-ALPPL2 antibody. Both the mesothelioma cell line M28 and the control BPH-1 line (nonbinding by M25) were used in the study. As shown in Figure 1C, anti-ALPPL2 antibody detected a 72 kDa band only in M28 but not BPH-1 IP products.

Finally, we performed ectopic expression and flow cytometry analysis to confirm target identification. We transiently transfected CHO-K1 cells with an ALPPL2 expression plasmid. In parallel, we produced recombinant M25 human IgG1. We analyzed M25 IgG1 binding to CHO-K1-ALPPL2 cells by flow cytometry. As shown in Figure 1D, M25 IgG1 binds specifically to CHO-K1-ALPPL2 but not CHO-K1, further validating that the target for M25 is ALPPL2.

### ALPPL2 is expressed by all subtypes of mesothelioma

We have previously reported IHC studies of the panel of anti-mesothelioma scFvs including M25 on frozen malignant mesothelioma tissues where we found positive staining in epithelioid, sarcomatoid, and mixed subtypes (An et al., 2008). To broaden applicability and examine a greater number of patient samples, we performed additional IHC studies on formalin fixed paraffin embedded (FFPE) mesothelioma tissues. The M25 antibody, like other antibodies selected from phage libraries on live tumor cells, binds to a conformational epitope and does not work on FFPE tissues (An et al., 2008; Bidlingmaier et al., 2009). As such, we performed IHC using two commercially available FFPE-compatible anti-ALPPL2 antibodies. In the first experiment, using an affinity purified anti-ALPPL2 rabbit antibody on human mesothelioma arrays containing a total of 131 tissue cores, about 72% of malignant mesothelioma tissues showed positive staining for ALPPL2 (33.6% strong, 19.1% moderate, and 19.1% weak) (Figure 1E and Supplemental Table S1). Normal adjacent tissues and infiltrating lymphocytes showed no detectable staining (Figure 1E). In a second experiment using a different FFPE-compatible mouse monoclonal antibody on a mesothelioma tissue array containing 100 cores, about 47% of mesothelioma tissues are stained positive (Supplemental Table S1). Positive staining was seen on epithelioid, sarcomatous (spindle) and mixed types. Taken together, although the exact percentage varies depending on the antibodies used and a subjective call for intensity cut off, it is evident that ALPPL2 is expressed by a significant portion of mesothelioma cases across subtypes.

### ALPPL2 expression shows exquisite tissue specificity

ALPPL2 expression in normal FFPE tissue was evaluated by using the FFPE-compatible anti-ALPPL2 rabbit antibody. A FFPE human normal tissue array that contains 66 cores was stained. Strong positive signal was observed only in placental trophoblasts. No staining was observed in any of the other normal tissues (Supplemental Table S2).

### Affinity maturation of M25 scFv

ALPPL2 represents a rare set of cell surface antigens that can be classified as being truly tumor specific due to its virtual non-expression on normal tissues as described above. It is an excellent target for the development of various types of therapies based on scFv, IgG, bispecific antibody, or scFv-targeted engineered T cells. We first measured M25 scFv binding to HEK293 cells expressing ALPPL2 by flow cytometry and obtained an apparent affinity value of 83 nM (Supplemental Figure S1). While this apparent affinity is sufficient for some therapeutic applications, we sought to broaden applicability and create a panel of affinity variants by yeast surface display-based affinity maturation (VanAntwerp and Wittrup, 2000). We first created yeast-displayed M25 scFv mutagenesis libraries (Bidlingmaier et al., 2015). We then selected high affinity binders to human ALPPL2 by FACS and identified the top two clones (M25FYIA and M25ADLF) that showed over 100-fold improvement in binding affinity as scFv (Supplemental Table S3). Importantly, like the parental M25 scFv, none of the high affinity variants bind to the closely related ALPI or the non-tissue specific ALPL (Supplemental Table S3). We thus have created a series of M25 variants with high specificity and a range of affinities, which can meet the demand of developing various types of therapies in the future using different antibody forms, i.e., scFv for chimeric antigen receptor T cell (CAR-T), or IgG for antibody-drug conjugate, or a combination of forms for bispecific.

### Evaluation of M25 IgG affinity and specificity

For this study, we are focused on studying therapeutic potential of ALPPL2 targeting using antibody-drug conjugates, and are thus focused on evaluating the IgG1 form of M25. To assess the affinity and specificity of the recombinant M25 IgG1 that we produced, a panel of tumor and normal cell lines were used for binding study by flow cytometry. As shown in Figure 2A, M25 IgG1 showed specific binding to the two mesothelioma cell lines studied, M28 (apparent K_D_ = 0.44 nM) and VAMT-1 (apparent K_D_ = 0.8 nM), but not to any of the normal cell lines studied, including human fibroblast cell line HS27, kidney cell line HK-2, and primary liver cell line HS775Li.

**Figure 2.**
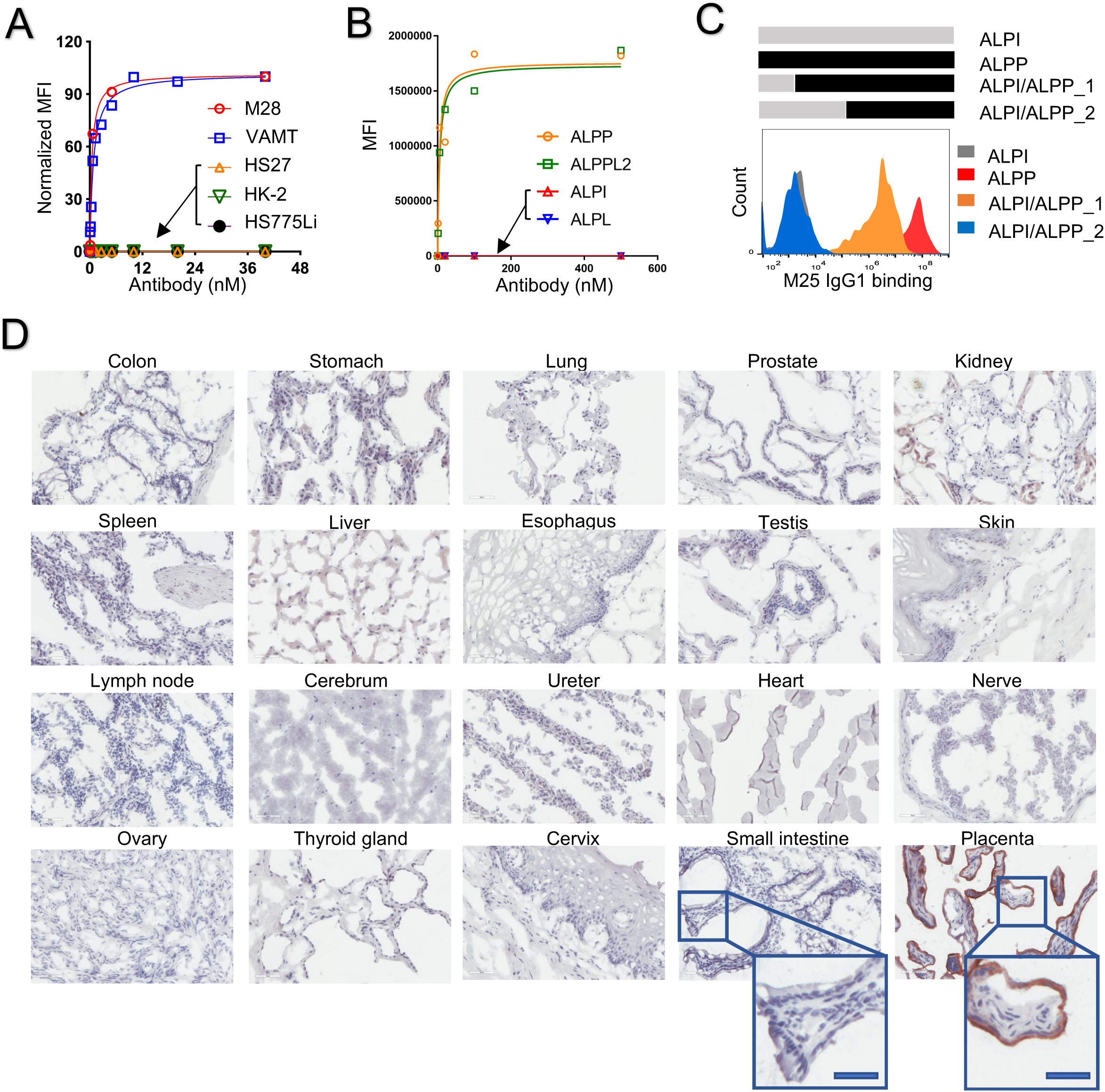
Specificity of M25 IgG binding to tumor cells and to different members of the alkaline phosphatase family. (A) M25 IgG1 binds specifically to mesothelioma cell lines M28 and VAMT-1 with no binding to control normal human cells (HS27, a normal foreskin fibroblast cell line; HK-2, a normal kidney epithelial cell line; and HS775Li, a normal primary liver cell line). Apparent K_D_ was estimated by curve-fitting (GraphPad). (B) M25 IgG1 binds specifically to ALPPL2 and ALPP, but not ALPI or ALPL. Flow cytometry analysis of M25 IgG1 binding to CHO-K1 cells transiently expressing each of the four members of the human alkaline phosphate family. ALPPL2, placental like-2 alkaline phosphatase; ALPP, placental alkaline phosphatase; ALPI, intestinal alkaline phosphatase; ALPL, tissue-nonspecific alkaline phosphatase. (C) Upper panel: Binding region determination using hybrid ALPI/ALPP fusion constructs. Sequence information is shown in Supplemental Figure 2S. Lower panel: flow cytometry analysis of binding to HEK293A cells expressing hybrid constructs. Red, binding to HEK293A cells expressing full-length ALPP. Grey, binding to HEK293A cells expressing full-length ALPI. Orange, binding to HEK293A cells expressing ALPI/ALPP hybrid_1. Blue, binding to HEK293A cells expressing ALPI/ALPP hybrid_2. (D) M25 IgG1 IHC staining of FDA standard panel of human frozen normal tissues. Other than placenta trophoblasts, none of the normal tissues studied showed positive staining. Scale bar: 50 μm. Enlarged windows are close-up views; scale bar (blue): 30 μm.

We further studied M25 IgG1 specificity towards each of the four members of the human alkaline phosphatase family: placental (ALPP), placental-like (ALPPL2), intestinal (ALPI), and liver/bone/kidney (ALPL, tissue non-specific). We studied binding specificity using CHO-K1 cells transfected with plasmids expressing the four genes, and found that M25 IgG1 binds only to ALPPL2 and ALPP but not the closely related ALPI and the tissue non-specific ALPL (Figure 2B). The apparent binding affinities of the M25 IgG1 for ALPPL2 and ALPP are about 0.5 nM on living cells, well suited for therapeutic development.

### Preliminary epitope mapping

As mentioned above, M25 was originally identified from phage antibody display library selection on live tumor cells and tissues (An et al., 2008), and does not bind to denatured proteins, thus recognizing a conformational, not linear, epitope. To perform a preliminary assessment of the M25 binding epitope, we created a series of hybrid molecules between ALPP (binding) and ALPI (non-binding, Supplemental Figure S2), expressed them in HEK293 cells and measured M25 IgG1 binding by flow cytometry (Figure 2C). The result suggests that M25 binds to a conformational epitope within amino acids 55-262 of ALPP.

### Examine tissue specificity of M25 IgG on a broad panel of frozen normal human tissues

To comprehensively study the tissue specificity of M25 IgG1, we performed IHC on a set of FDA standard frozen normal human tissue arrays that cover 30 different organs/tissues in triplicate (90 tissue cores total). Positive staining was only observed in placental trophoblasts, and no staining was seen for any other normal organ/tissue (Figure 2D and Table 1, with additional studies shown in Supplemental Figure S3), demonstrating an exquisite specificity. The same tissue specificity was also observed for the high affinity variant M25FYIA IgG1 (Supplemental Table S4). Those IHC results on frozen human tissues, together with the results from FFPE tissues described above, validate tissue specificity of ALPPL2 expression and targeting specificity of M25 IgG1.

**Table 1.**
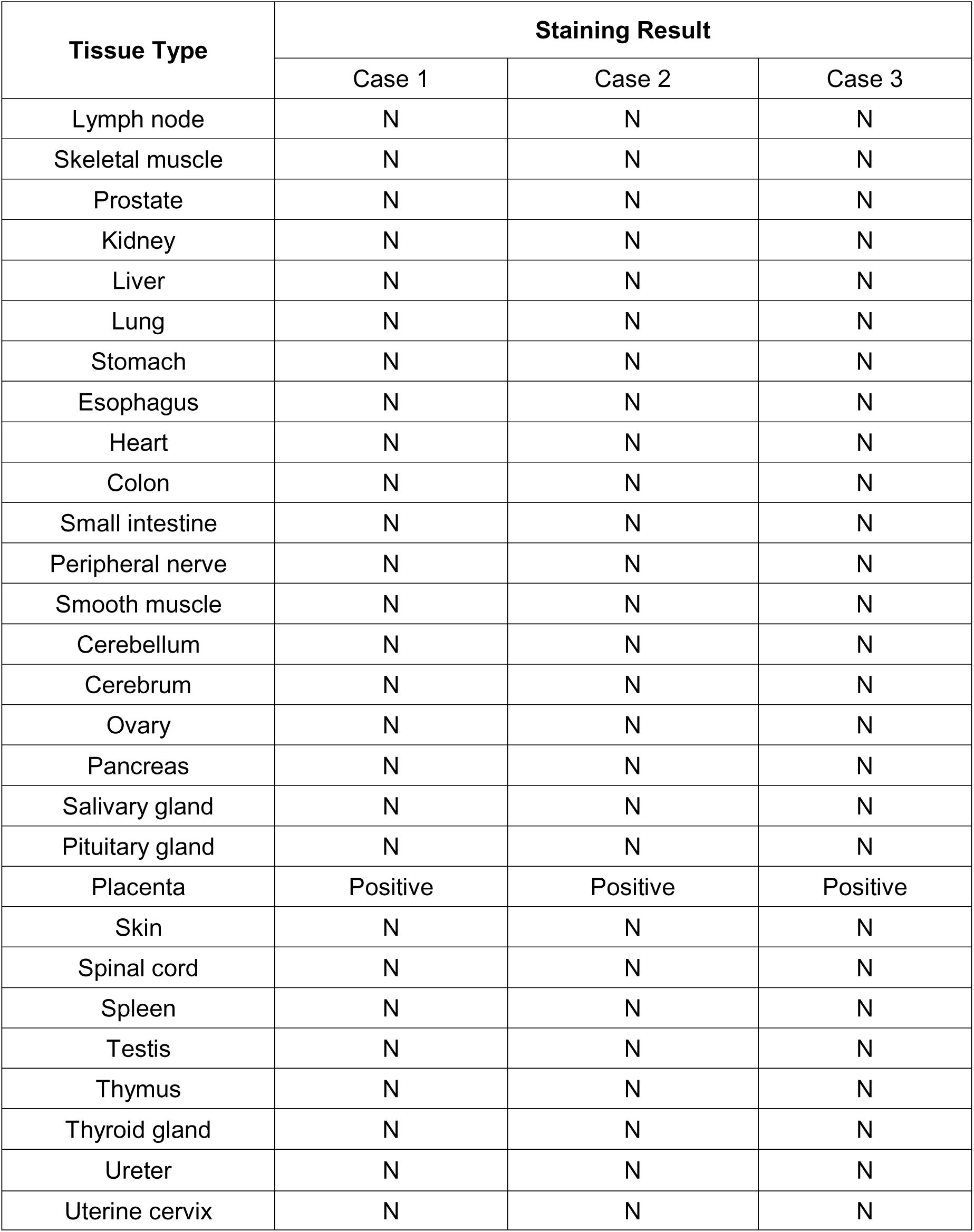
Summary of IHC staining patterns of M25 IgG1 on a frozen normal human tissue array. N: No staining

### Evaluation of the anti-ALPPL2 antibody M25 as a potential tumor-specific delivery vehicle

We studied M25 internalization on mesothelioma cell lines M28 (epithelial type) and VAMT-1 (sarcomatous type) by fluorescence microscopy. As shown in Figure S4A, M25 IgG1 was internalized by both M28 and VAMT-1 cells at a moderate rate, with clear evidence of internalization occurring after 24h.

We next studied M25 internalization using a functional internalization assay, which assesses the ability of M25 to mediate intracellular delivery of a plant toxin (saporin) that does not enter cells on its own (Stirpe et al., 1992; Vago et al., 2005). Potent and selective antitumor activity was observed against both M28 and VAMT-1 cells but not any of the control cells studied (HS27, HK-2 and HS775Li) (Supplemental Figure S4B). These data suggest that our anti-ALPPL2 antibody M25 is suitable for constructing targeted therapeutics that require intracellular payload delivery.

### Anti-ALPPL2 ADC development

We next created an anti-ALPPL2 ADC by conjugating either of the two microtubule inhibitors, monomethyl auristatin F (MMAF) (Doronina et al., 2006) or monomethyl auristatin E (MMAE) (Senter and Sievers, 2012), to the M25 IgG1 via a protease-cleavable linker maleimidocaproyl-valine-citrulline-p-aminobenzoyloxycarbonyl (MC-vc-PAB) (Sherbenou et al., 2016; Su et al., 2018). The drug to antibody ratio (DAR) was determined by hydrophobic interaction chromatography (HIC) (Sherbenou et al., 2016; Su et al., 2018). As shown in Figure 3A and 3B, the DAR of M25-MCvcPAB-MMAF and M25-MCvcPAB-MMAE were 3.3 and 3.4, respectively. A non-binding human IgG1 was also conjugated to both drugs (Ctrl IgG-MCvcPAB-MMAF and Ctrl IgG-MCvcPAB-MMAE) with similar DARs (Supplemental Figure S5A and S5B). Conjugation of MMAF and MMAE did not alter the binding properties (affinity and specificity) of the M25 IgG (Supplemental Figure S5C).

**Figure 3.**
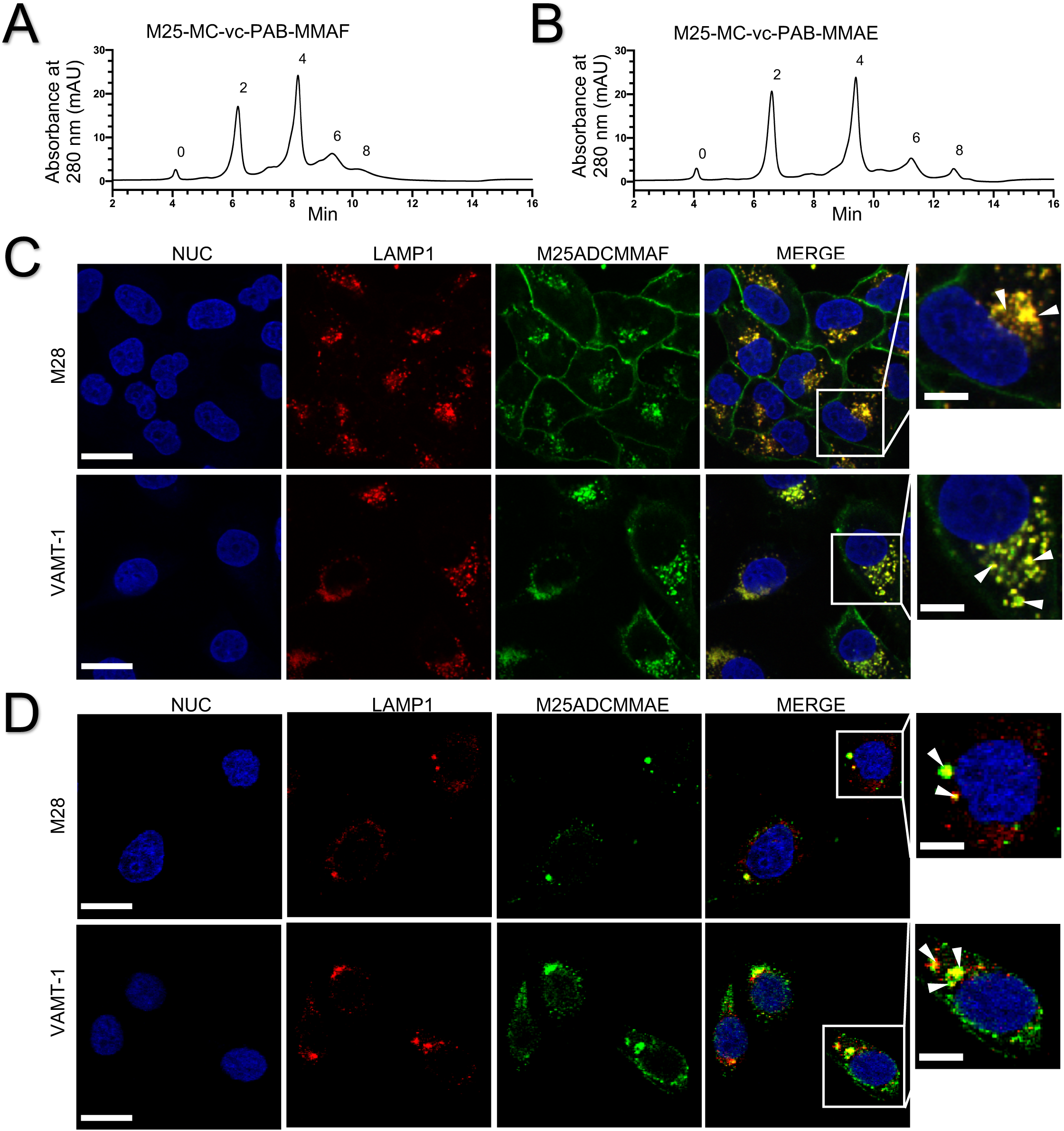
Internalization of anti-ALPPL2 ADC into mesothelioma cells. (A) Generation and HPLC analysis of anti-ALPPL2 ADCs. MMAF-conjugated M25 IgG1 (M25-MC-vc-PAB-MMAF or M25ADCMMAF) (A) and MMAE-conjugated M25 IgG1 (M25-MC-vc-PAB-MMAE or M25ADCMMAE) (B) were analyzed by HIC. (C) Confocal microscopy study of internalization into mesothelioma cell lines (M28 and VAMT-1) by M25ADCMMAF. (D) Confocal microscopy study of internalization into mesothelioma cell lines by M25ADCMMAE (D). For both (C) and (D), co-localization (yellow) between the lysosomal marker LAMP1 (red) and ADC (green) after 4h of incubation are seen in a single confocal slice. Nuc, nuclei. Scale bar: 30 μm. Enlarged windows on the far right show close-up views of co-localization (yellow dots pointed by white arrowheads); scale bar: 10 μm.

### Internalization of anti-ALPPL2 ADC

Internalization of MMAF and MMAE-conjugated M25 ADCs (M25ADCMMAF and M25ADCMMAE, respectively) was studied on mesothelioma cells by confocal fluorescence microscopy. On both M28 and VAMT-1 cells, M25ADCMMAF internalization was detected during the first 4h of incubation (Figure 3C) and continually increased over the 24h incubation period (Supplemental Figure S6). In contrast, the MMAE-conjugated ADC showed higher internalization activity with complete internalization of M25ADCMMAE detected during the first 4h of incubation (Figure 3D).

To further evaluate antibody localization to specific intracellular compartments, we studied trafficking post internalization of M25ADCMMAF and M25ADCMMAE using the lysosomal maker, lysosomal-associated membrane protein 1 (LAMP1). M25ADCMMAF colocalization with LAMP1 was observed in M28 and VAMT-1 mesothelioma cells after 4h of incubation (Figure 3C) and continually increased during the 24h incubation period (Supplemental Figure S6). M25ADCMMAE colocalization with LAMP1 was observed in M28 and VAMT-1 cells after 4h of incubation (Figure 3D). Thus M25 ADC is internalized and trafficked into the lysosome, a preferred subcellular location for ADC where degradation by lysosomal proteases releases the small molecule drug from the antibody to exert its therapeutic effect (de Goeij and Lambert, 2016).

### Anti-ALPPL2 ADC exhibits potent and specific tumor cytotoxicity in vitro

Anti-ALPPL2 ADC and control ADC were assessed for cytotoxicity *in vitro* against mesothelioma (M28 and VAMT-1) and control cells (normal human fibroblasts, kidney cells, and primary liver cells). As shown in Figure 4, anti-ALPPL2 ADCs potently reduced the viability of both mesothelioma cell lines but none of the control cells. The EC50 value was estimated to be 0.54 nM for M25ADCMMAF on M28, and 67 nM on VAMT-1; and 0.03 nM for M25ADCMMAE on M28, and 0.44 nM on VAMT-1. In contrast, > 1,000-fold higher concentrations were required for control ADCs to have similar effects. No cytotoxic effect was observed on control normal cells for either M25ADCMMAF or M25ADCMMAE.

**Figure 4.**
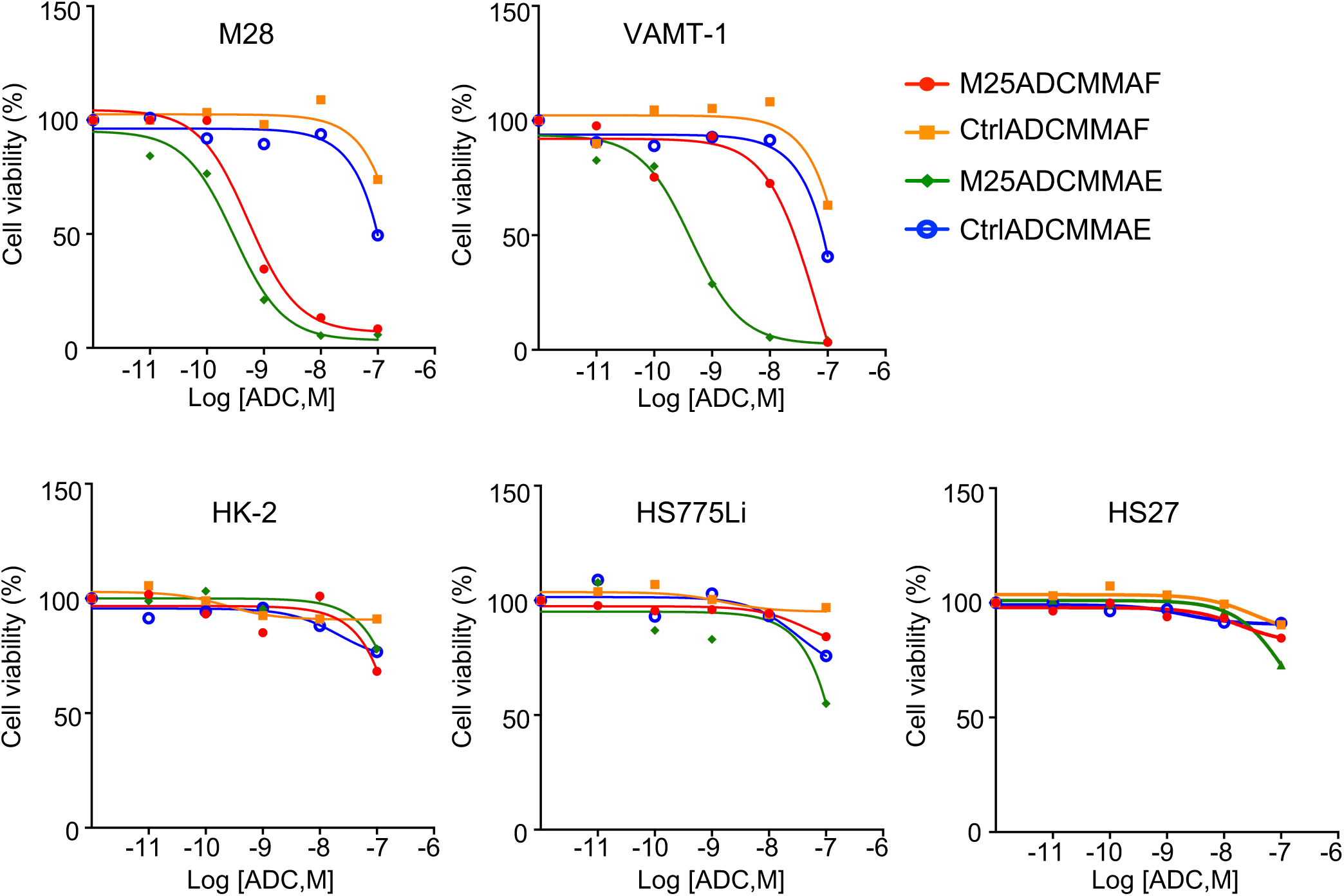
Tumor-specific cytotoxicity of anti-ALPPL2 ADCs *in vitro*. M25-MC-vc-PAB-MMAF, M25-MC-vc-PAB-MMAE, Ctrl IgG-MC-vc-PAB-MMAF, and Ctrl IgG-MC-vc-PAB-MMAE were incubated with mesothelioma cell lines (M28 and VAMT-1) and control normal human cell lines (HS27, HK-2, and HS775Li). Normalized cell viability is shown as a function of the ADC concentration. M25 ADCs showed potent cytotoxic effects on mesothelioma cell lines but not any of the control normal human cells lines. EC50 values were estimated by curve fitting (Prism, GraphPad): M25ADCMMAF, 0.54 nM on M28 and 67 nM on VAMT-1; M25ADCMMAE, 0.03 nM on M28 and 0.44 nM on VAMT-1.

### Anti-ALPPL2 ADC shows curative potential in mesothelioma xenograft mouse models

The *in vivo* efficacy of anti-ALPPL2 ADC was evaluated in mice carrying mesothelioma cell line xenografts. Both the epithelial (M28) and sarcomatous (VAMT-1) xenografts were studied. An outline of the study design is shown in Figure 5A. We studied both the MMAF- and the MMAE-based ADCs. First, nude mice were subcutaneously (s.c.) implanted with one million M28 cells. Tumor growth was palpated, and the sizes were measured by caliper. When the average tumor volume reached 250 mm^3^, the mice were randomized into three study groups and treated intravenously (i.v.) with M25ADCMMAF, naked M25IgG, or vehicle control (PBS) on Study Day 0 at 5 mg/kg every 4 days for a total of 4 injections (Q4D×4). As shown in Figure 5B, tumor growth is inhibited by M25ADCMMAF but not vehicle control or naked antibody. No body weight loss was observed in all treatment groups (Supplemental Figure S7A).

**Figure 5.**
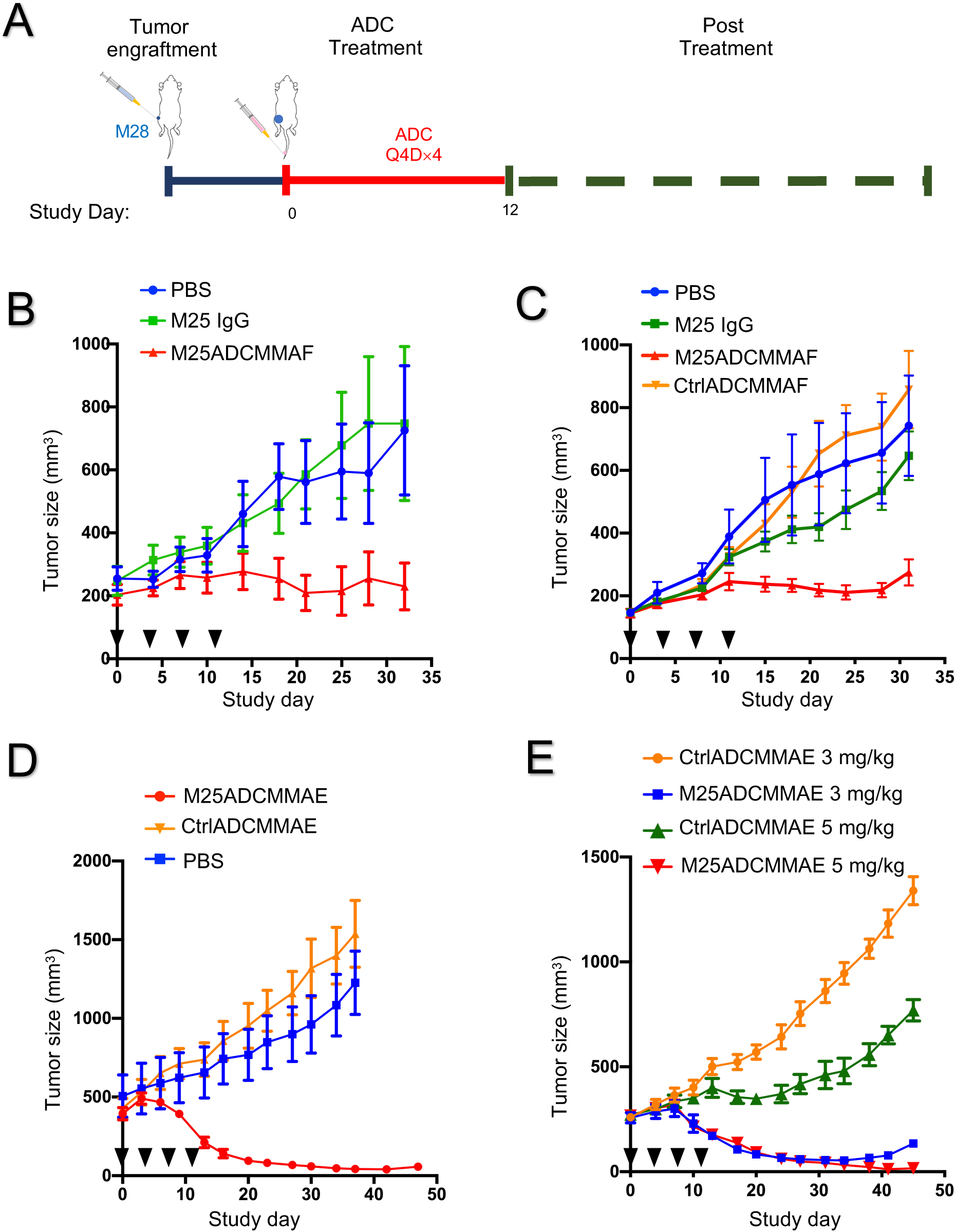
Anti-tumor activity of anti-ALPPL2 ADCs (M25-MC-vc-PAB-MMAF or M25ADCMMAF; and M25-MC-vc-PAB-MMAE or M25ADCMMAE) *in vivo*. (A) Outline of the experimental design. (B) Nude mice bearing M28 xenografts (∼250 mm^3^) were treated with M25ADCMMAF for a total of 4 doses at 5 mg/kg, along with the two control arms, vehicle only (PBS) and naked antibody (M25 IgG1). Injection days are indicated by black triangles. (C) NSG mice bearing M28 xenograft (∼150 mm^3^) were treated with M25ADCMMAF, CtrlADCMMAF (Ctrl IgG-MC-vc-PAB-MMAF), vehicle only (PBS) and naked antibody (M25 IgG1) at 5 mg/kg for a total of 4 doses. Injection days are indicated by black triangles. Quantification of tumor sizes shows a significant difference (* p < 0.05) at the end of the experiment. Student’s t test, unpaired two-tailed. (D) NSG mice bearing large-sized (∼ 500 mm^3^) M28 xenografts were i.v. injected with M25ADCMMAE, CtrlADCMMAE and vehicle only (PBS) at 5 mg/kg for a total of 4 doses. Injection days are indicated by black triangles. Near complete regression of M28 tumors was observed. (E) NSG mice bearing M28 xenograft (∼ 250 mm^3^) were treated with M25ADCMMAE or CtrlADCMMAE at 5 mg/kg or 3 mg/kg very four days for a total of 5 doses. Injection days are indicated by black triangles. Complete eradication of M28 tumors was observed in the 5 mg/kg group.

We performed an additional study in NSG mice carrying M28 cell line xenografts. When the average tumor volume reached 150 mm^3^, the mice were randomized into four study groups and i.v. injected (Q4D×4) with M25ADCMMAF, CtrlADCMMAF, naked M25 IgG1 or PBS on Study Day 0 at 5 mg/kg. Tumor growth was inhibited by M25ADCMMAF but not vehicle control (PBS), naked IgG1, or the control ADC (Figure 5C). No body weight loss was observed for all treatment groups (Supplemental Figure S7B).

We next studied the efficacy of MMAE-based ADCs. In the first experiment, M28 cells were s.c. implanted into NSG mice and allowed to grow to ∼500 mm^3^ before ADC treatment (5 mg/kg, Q4D×4). The M25ADCMMAE was able to greatly reduce the size of this large tumor (> 90% reduction) (Figure 5D). Neither vehicle nor the CtrlADCMMAE affected tumor growth. No body weight loss was observed in all treatment groups (Supplemental Figure S8).

To further evaluate in vivo efficacy of M25ADCMMAE, we tested two different ADC doses in NSG mice carrying M28 xenografts. When tumors reached 250 mm^3^, M25ADCMMAE and CtrlADCMMAE at 3 mg/kg or 5 mg/kg were i.v. injected every 4 days for a total of 5 times (Q4D×5). Both 3 mg/kg and 5 mg/kg M25ADCMMAE resulted in potent tumor size reduction. Complete tumor eradication was observed in the 5 mg/kg dosing group (Figure 5E). CtrlADCMMAE did not have any significant effect on tumor growth (Supplemental Figure S9). No body weight loss was observed in all treatment groups (Supplemental Figure S10).

To evaluate M25ADCMMAE simultaneously on both types of mesothelioma *in vivo*, we generated a dual xenograft model where NSG mice were s.c. grafted with both M28 (right flank) and VAMT-1 (left flank) cell lines. M28 tumors grew rapidly to reach an average size of 500 mm^3^ while VAMT-1 tumors reached an average size of 150 mm^3^.

M25ADCMMAE and CtrlADCMMAE were i.v. injected weekly for a total of 4 times (Q7D×4) at 5 mg/kg (Figure 6A). As shown in Figure 6B-D, both M28 and VAMT-1 xenografts responded to M25ADCMMAE treatment. The size of M28 tumors (> 500 mm^3^) was reduced by 90-100% on Day 45 (absolute tumor size shown in Figure 6B with normalized size shown in Figure 6D) while VAMT-1 was not detectable by caliper measurement on Day 51 (absolute tumor size shown in Figure 6B with normalized size shown in Figure 6C). No body weight loss was observed in all treatment groups (Supplemental Figure S11).

**Figure 6.**
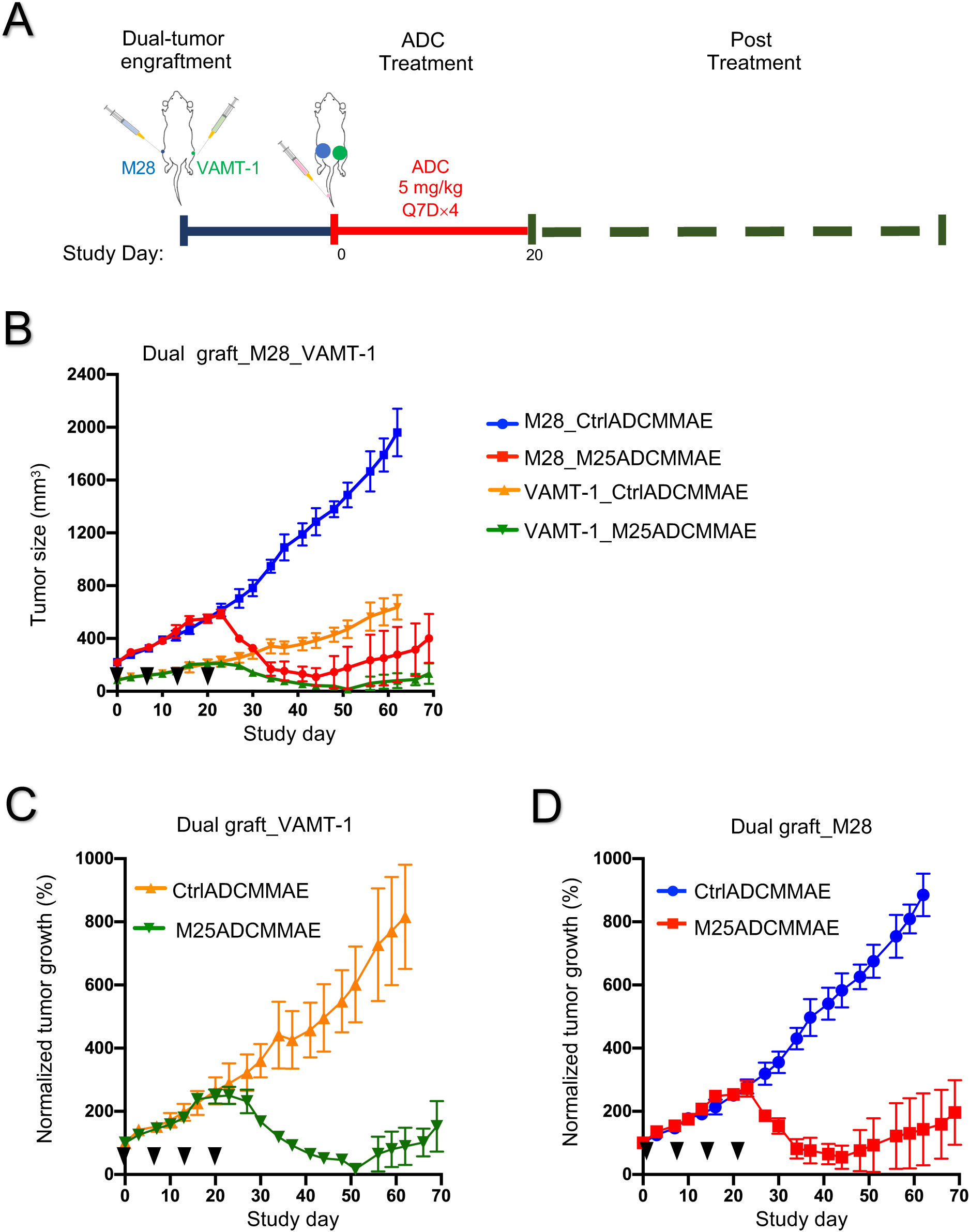
In vivo anti-tumor activity of anti-ALPPL2 ADC (M25ADCMMAE) in a dual tumor xenografts model. (A) Experimental scheme. (B) NSG mice bearing dually grafted tumor xenografts, M28 (right flank, ∼500 mm^3^) and VAMT-1 (left flank, ∼150 mm^3^), were i.v. injected with M25ADCMMAE and CtrlADCMMAE at 5 mg/kg weekly for total of 4 doses. (C) Normalized tumor growth of VAMT-1 xenografts. (D) Normalized tumor growth of M28 xenografts.

Taken together, ALPPL2-targeted ADCs constructed from the M25 IgG1 are efficacious in both types of mesothelioma xenograft models, with ADC conjugated with MMAE being more potent and showing curative potential.

## Discussion

Target identification is a major challenge in cancer therapy development. The vast majority of tumor-associated antigens are not tumor specific, as their expression can be found on multiple normal tissues. A specific challenge in mesothelioma therapy development is the lack of tumor-specific cell surface antigens that are expressed by all subtypes of the disease but not normal tissues. Mesothelin is the most frequently targeted cell surface antigen for mesothelioma. However, while mesothelin is expressed by epithelial mesothelioma, it is rarely expressed by the difficult-to-treat sarcomatous mesothelioma. In addition, mesothelin is also expressed by normal mesothelium, raising the issue of on-target toxicity even for therapies based on local delivery.

To meet this challenge, we have taken an unbiased approach and selected phage antibody libraries on live mesothelioma tissues and cells to identify novel antibodies that bind to all subtypes of mesothelioma but not normal mesothelium and other normal tissues (An et al., 2008). The approach is entirely unbiased with no prior knowledge of the target antigen. As reported previously (An et al., 2008), we identified a panel of scFvs that bind to both epithelial and sarcomatous mesothelioma, but not the normal mesothelium. Among these scFvs, M25 shows the highest specificity (An et al., 2008) and in this study we have identified the target antigen bound by M25 as ALPPL2, a member of the human alkaline phosphatase family. We show that our M25 antibody binds specifically to ALPPL2 and ALPP, both of which are exclusively expressed by placental trophoblasts, but not ALPI and ALPL that are expressed on normal tissues. Using FDA standard panel of frozen human tissue arrays, we confirm that M25 binds to placenta trophoblasts but none of the 29 other normal human tissues, showing an exquisite tissue specificity that should enable the development of various anti-cancer therapies. To evaluate therapeutic potential of targeting this novel tumor antigen, we constructed M25 ADCs conjugated with either MMAF or MMAE, and showed that M25 ADCs kill both epithelial and sarcomatous mesothelial cell lines in vitro and in mouse xenograft models, with MMAE-based ADC showing higher potency *in vivo*.

Although the biological function of ALPPL2 in tumor remains unclear, it is, by all practical measures, a true tumor specific antigen. Other than placental trophoblasts, ALPPL2 expression on normal tissues is virtually absent, providing an excellent opportunity for developing therapies that require a high degree of tumor specificity. Besides microtubule inhibitors (MMAE and MMAF) as we showed in this study, anti-ALPPL2 antibodies can be conjugated with other classes of drugs such as DNA crosslinking agents or radionuclides (e.g., alpha particles). In addition, anti-ALPPL2 antibodies can be used for construction of bispecific T cell engagers, CAR-T cells and immunocytokines.

Although this study is focused on mesothelioma, we have data suggesting that ALPPL2 is also expressed by several other tumor types such as testicular, ovarian, pancreatic, gastric, colorectal, and lung cancers (Yang Su and Bin Liu, unpublished observations), consistent with reports in literature regarding this target (e.g., seminoma (Koshida and Wahren, 1990; Lange et al., 1982), ovarian cancer (Haije et al., 1979; Orsaria et al., 2016), pancreatic cancer (Dua et al., 2013), gastric cancer (Liu et al., 2019), colorectal cancer (Skinner and Whitehead, 1981), and lung cancer (Wick et al., 1987)). Testicular cancer showed virtually 100% positive ALPPL2 expression by IHC. The other tumors mentioned above showed positive IHC staining in a fraction of cases (Yang Su and Bin Liu, unpublished observations). For clinical application to those tumors, future biomarker development is desired to aid patient stratification and identify those who are most likely to respond.

There have been a couple of previous reports on generation of ALPPL2 or ALPP targeting agents. An anti-ALPPL2 aptamer was identified in a study on pancreatic cancer (Dua et al., 2013). It is unclear, however, if this aptamer is specific to ALPPL2 or ALPP and if it binds to the closely related ALPI and the tissue non-specific ALPL. In a second study using phage display panning on recombinant proteins, an anti-ALPP scFv was identified and studied *in vivo* for tumor imaging (Ravenni et al., 2014). This scFv, however, does not bind to ALPPL2 (Ravenni et al., 2014). In contrast, our scFv was isolated by panning on live tumor tissues/cells and recognizes both ALPP and ALPPL2 but none of the other two members of the alkaline phosphatase family, thus differentiating from previously generated agents.

In summary, we have identified the target antigen of a tumor specific antibody M25 as ALPPL2, a member of the human alkaline phosphatase family that is expressed by placental trophoblasts. We confirmed the exquisite tissue specificity of M25 on normal human tissues. We showed that > 47% of mesothelioma expresses ALPPL2. We showed that M25 ADCs are effective against both epithelial and sarcomatous mesothelioma cell lines *in vitro* and *in vivo*. Besides ADC, this highly specific antibody-antigen pair lends itself to additional translational development including bispecific and CAR-T against ALPPL2 expressing tumors that are resistant to current treatments.

## Materials and Methods

### Cell Culture

Human mesothelioma cell lines VAMT-1 and M28 were originally obtained from Dr. Brenda Gerwin‘s lab at the National Cancer Institute, and maintained in the lab in RPMI1640 supplemented with 10% fetal bovine serum (FBS) and 100 μg/ml penicillin-streptomycin (Thermo Fisher Scientific). Human foreskin fibroblast cell line HS27 and kidney cell line HK-2 were obtained from American Type Culture Collection (ATCC) and cultured according to supplier’s instruction. HEK293A cell line is purchased from Sigma-Aldrich, and cultured according to supplier’s instruction. Primary normal human liver cell line HS775li was obtained from UCSF Cell Culture facility. The benign prostatic hyperplasia epithelial cell line BPH-1 was originally obtained from Dr. Jerry Cunha’s lab at UCSF and maintained in DMEM supplemented with 10% FBS and 100 μg/ml penicillin-streptomycin. All cell lines were cultured in humidified atmosphere of 95% air and 5% CO_2_ at 37°C.

### Recombinant M25 scFv-Fc and IgG1

To generate M25 scFv-Fc and M25 full-length human IgG1, M25 scFv was subcloned into either the Fc fusion expression vector pFUSE-hIgG1 Fc2 (InvivoGen) or a modified Abvec expression vector that was originally provided by Dr. Patrick Wilson (Smith et al., 2009), respectively. The antibodies were transiently expressed in HEK 293A cells following polyethylenimine (Sigma-Aldrich) transfection (Sherbenou et al., 2016; Su et al., 2018). The secreted antibody in serum-free DMEM containing Nutridoma-SP (Sigma) and penicillin-streptomycin were harvested 5 days post transfection, filtered by 0.22 μm filter (Millipore), and purified using protein A agarose (Pierce/Thermo Fisher Scientific). Antibody concentrations were determined using nanodrop 2000 (Thermo Fisher Scientific).

### Immunoprecipitation and mass spectrometry

M28 and BPH-1 cells were surface biotin-labeled with EZ-Link Sulfo-NHS-LC-Biotinylation Kit (Pierce/Thermo Fisher Scientific) and lysed with cell lysis buffer (20 mM Tris-HCl, pH 7.4, 0.3 M NaCl, 1% Nonidet P-40) supplemented with complete protease inhibitor cocktail (Roche). The M25 scFv-Fc was immobilized onto protein A beads (Thermo Fisher Scientific) and cross-linked by dimethyl pimelimidate dihydrochloride (DMP) (final concentration at 20 mM) (Sigma) at RT for 30 min. The reaction was stopped by incubation in ethanolamine (0.2M, pH8.0) at RT for 2h. The beads were washed with PBS and incubated with biotin-labeled cell lysates at 4°C overnight, washed with 0.1%PBS/Tween20 containing 500 nM NaCl followed by additional wash (> 3 times) with PBS. The beads with antigens captured were loaded onto a SDS-PAGE gel (4-20% gradient polyacrylamide) (Thermo Fisher Scientific) in duplicates, one for Gelcode staining (Thermo Fisher Scientific), and the other Western blot analysis to locate the membrane protein band by streptavidin-conjugated with horseradish peroxidase (HRP) (Jackson ImmunoResearch Laboratories) and ECL substrate (Thermo Fisher Scientific). Bands on the Gelcode-stained gel corresponding to positions on the Western blot were excised, in-gel digested with trypsin and analyzed by tandem mass spectrometry (MS/MS, Mass Spectrometry Facility, University of Minnesota). Spectra were searched using Protein Prospector against the SwissProt database (Ha et al., 2014).

### Immunohistochemistry

IHC studies on frozen and FFPE tissues were performed as described (Su et al., 2018). For IHC on frozen normal human tissue arrays (US Biomax), slides were air-dried for 20 min at RT, fixed in 4% paraformaldehyde for 15 min, washed 3 times with PBS, and incubated with 3% H_2_O_2_ (Thermo Fisher Scientific) for 10 min to block endogenous peroxidase activity. The slides were washed with PBS, further blocked with 2% donkey serum (Santa Cruz Biotechnology) and avidin/biotin (Vector Laboratories), washed twice with PBS, incubated with 15 μg/ml biotinylated M25 IgG1 at 4 C overnight, washed three times with PBS, further incubated with streptavidin-HRP (Jackson ImmunoResearch Laboratories) followed by detection using diaminobenzidine as substrate (Pierce/Thermo Fisher Scientific).

FFPE tissue arrays (US Biomax) were deparaffinized in xylene overnight and rehydrated by sequential exposure to 100%, 95%, 70% ethanol, and ddH_2_O (5 min each). Slides were treated with Tris-EDTA Buffer (10 mM Tris Base, 1 mM EDTA Solution, 0.05% Tween 20, pH 9.0) at 95°C for 20 min followed by PBS wash. Endogenous peroxidase activity was blocked using 3% H_2_O_2_ for 15 min followed by PBS wash. After blocking with 2% donkey serum at RT for 1h, anti-ALPPL2 affinity-purified rabbit polyclonal antibody (Origene) or anti-ALPPL2 mouse antibody (Clone SPM593, Lifespan Biosciences) were diluted 1:500 into 2% donkey serum/PBS and incubated with the tissue slides at 4°C overnight. After PBS wash, slides were incubated with anti-rabbit or anti-mouse DAKO EnVision™+, and detected by liquid DAB+ (DAKO). Slides were counterstained with hematoxylin followed by the bluing reagent (Scytek). Slides were mounted in aqua-mount (Lerner Laboratories), and scanned by Aperio Digital Pathology Scanner (Leica Biosystems).

### Specificity assessment by ectopic expression and flow cytometry

CHO-K1 or HEK293 cells were transfected with plasmids expressing human ALPPL2, ALPP, ALPI, and ALPL using Lipofectamine® 2000 (Life Technologies/Thermo Fisher Scientific), and incubated with anti-ALPPL2 human antibodies followed by Alexa Fluor® 647-conjugated goat anti-human IgG (Jackson ImmunoResearch Laboratories). Binding was analyzed by flow cytometry (BD Accuri™ C6, BD Biosciences) that generated median fluorescence intensity (MFI) values.

### Immunotoxin assay to assess functional internalization

Immunotoxin study was performed as described (Su et al., 2018). Briefly, mesothelioma (M28 and VAMT-1) and control (HS27, HS775li, and HK-2) cells were seeded at 1,000 cell/well in 96-well plate (Falcon) and cultured in a 5% CO_2_ incubator at 37°C overnight. Biotinylated M25 IgG1 and Streptavidin-ZAP (Advanced Targeting Systems) were mixed at a molar ratio of 1:1 on ice for 30 min and added to cells at final concentration of SA-ZAP at 0, 0.6, 6, 60, 300 nM at 37°C for 96h. Cell viability was determined using a CCK-8 assay kit following the manufacturer’s instructions (Dojindo).

### Apparent cell binding affinity

Apparent K_D_ was measured on live cells using methods described previously (Sherbenou et al., 2016; Su et al., 2018). Briefly, mesothelioma and control cells were grown to exponential phase, detached by trypsin digestion and suspended in ice-cold PBS buffer with 2% FBS. M25 IgG was serially diluted into binding buffer and incubated with 1× 10^5^ cells in duplicated wells overnight at 4°C. Cells were washed three times with ice-cold PBS and further incubated with Alexa Fluor® 647-conjugated goat anti-human IgG for 1h at 4°C. Binding was measured by flow cytometry, and apparent K_D_ was determined by curve fitting MFI values using Prism software (GraphPad).

### Antibody-drug conjugates

Antibody-drug conjugation was performed using methods described previously (Sherbenou et al., 2016; Su et al., 2018). Briefly, M25 IgG1 was partially reduced with 2 equivalents of Tris(2-carboxyethyl) phosphine hydrochloride (TCEP, Thermo Fisher Scientific) at 37 °C for 2h. The mixture was purified by Zeba spin column (Pierce/Thermo Fisher Scientific), and buffer-exchanged into PBS with 5 mM EDTA. Six equivalents of maleimidocaproyl valine-ciytulline-p-aminobenzoyloxycarbonyl-monomethyl auristatin phenylalanine (mcvcpabMMAF, ref. (Sherbenou et al., 2016; Su et al., 2018)) or maleimidocaproyl-valine-citrulline-p-aminobenzoyloxycarbonyl-monomethyl auristatin E (mcvcpabMMAE, Concortis) were incubated with TCEP-reduced M25 IgG at RT for 1h. Excess mcvcpabMMAF or mcvcpabMMAE was removed by running twice through the Zeba spin column. Purified ADCs were analyzed by HIC-HPLC with infinity 1220 LC System (Agilent). The drug-to-antibody ratio (DAR) is estimated from area integration using the OpenLab CDS software (Agilent).

### Immunofluorescence confocal microscopy

M28 and VAMT-1 cells (5,000 per well) were seeded in glass chamber slides (Thermo Fisher Scientific) and grown at 37°C overnight, incubated with 15 μg/ml M25 IgG or M25ADCs for 4h or 24h, washed with PBS, fixed with 4% formaldehyde, and permeabilized with PBS containing 0.1% Triton X-100 and 1% BSA. M25 IgG was detected by Alexa Fluor® 647-conjugated goat anti-Human IgG (Jackson ImmunoResearch Laboratories). Cell nuclei were counterstained with Hoechst (Life Technologies/Thermo Fisher Scientific). Lysosomes were marked by anti-LAMP1 antibodies (Clone D2D11, Cell Signaling Technology) followed by detection with FITC– labeled donkey anti-Rabbit IgG (Jackson ImmunoResearch Laboratories). Images were taken and analyzed by an automated FluoView confocal microscope (Olympus).

### *In vitro* ADC cytotoxicity assay

Mesothelioma (M28 and VAMT-1) and control (HS775li and HK-2) cells were seeded in 96-well plates at 2,000 cell/well and incubated with serially diluted ADCs at indicated concentrations for 96h at 37 °C in humidified chamber with 5% CO_2_. The media was then removed and cells gently washed once with PBS. 1 μM calcein AM (Invitrogen/Thermo Fisher Scientific) in PBS was added, and cells were incubated at RT for 40 min. Plates were then read on a fluorescence plate reader (Biotek) using an excitation wavelength of 485 nm and an emission wavelength of 530 nm. Percent of cell survival was normalized against mock-treated cells, and EC50 was determined by curve fitting using Prism software (GraphPad).

### *In vivo* efficacy study on mesothelioma subcu xenografts

All animal studies were approved by the UCSF Animal Care and Use Committee (AN092211) and conducted in adherence to the NIH Guide for the Care and Use of Laboratory Animals. Male and female NSG mice (NOD.Cg-Prkdc^scid^ Il2rg^tm1Wjl^/SzJ, The Jackson Laboratory) or nude mice (Ncr nude, sp/sp, Taconic) of 7 to 9 week of age were implanted with 1 × 10^6^ mesothelioma cells in 100 μL of 50% Matrigel subcutaneously at the flank of the animal. Growing tumors were palpated, and sizes measured by a caliper.

Tumor volume was calculated using the formula V = ½ (length x width^2^). When the tumor reached the desired size, animals were randomized into group of six and dosed intravenously with dosing schemes as indicated. Body weight and other signs of overt toxicity were monitored daily.

## Author contributions

YS, SB and BL designed experiments. YS, XZ, SB, CRB and NKL performed experiments. YS, SB and BL wrote the manuscript. BL conceived the overall project idea. All authors contributed to manuscript revision.

## Acknowledgment

We thank Dr. Yue Liu for advice on immunoprecipitation, Dr. Byron C. Hann for advice on animal study, Donghui Wang, Paul Phojanakong, Fernando Salangsang and Julia Malato at the UCSF Preclinical Therapeutic Core for performing animal studies, and the Center for Mass Spectrometry and Proteomics at the University of Minnesota for mass spectrometry analysis. This work is supported in part by the National Institutes of Health (R01 CA129491, CA118919, and CA171315 to BL).

