## Supplemental Tables + figures for "ALPPL2 is a highly specific and targetable tumor cell surface antigen"

**Supplemental Table S1.** ALPPL2 expression in mesothelioma tumor tissues.

**Supplemental Table S2.** ALPPL2 expression in normal tissues (IHC on FFPE tissue).

**Supplemental Table S3.** Affinity assessment of anti-ALPPL2 antibodies.

**Supplemental Table S4.** Summary of IHC staining patterns of M25 FYIA IgG1 on a frozen normal human tissue array.

**Supplemental Figure S1.** Apparent binding affinity of M25 scFv to cells expressing ALPPL2.

**Supplemental Figure S2.** Amino acid sequences of the two ALPI/ALPP fusion constructs.

**Supplemental Figure S3.** M25 IgG1 IHC staining of human frozen normal tissues.

**Supplemental Figure S4.** Tumor cell-specific internalization and cytotoxicity.

**Supplemental Figure S5.** Characterization of anti-ALPPL2 ADC.

**Supplemental Figure S6.** Subcellular localization of anti-ALPPL2 ADC in mesothelioma cells.

**Supplemental Figure S7.** Body weight assessment.

**Supplemental Figure S8.** Body weight assessment.

**Supplemental Figure S9.** Statistical analysis of tumor volumes.

**Supplemental Figure S10.** Body weight assessment.

**Supplemental Figure S11.** Body weight assessment of dually grafted mice.

**Supplemental Table S1**. ALPPL2 expression in mesothelioma tumor tissues. IHC study using two different antibodies was performed on FFPE tumor arrays. The number and percentage of each type of staining patterns are indicated. +++: Strong positive staining. ++: Moderately positive staining. +: Weak positive staining. -: No staining.

| **Staining Result** | **Antibody 1 (Mouse mAb)** | **Antibody 2 (Rabbit pAb)** |
|---|---|---|
| +++ | 20 (20%) | 44 (33.6%) |
| ++ | 14 (14%) | 25 (19.1%) |
| + | 13 (13%) | 25 (19.1%) |
| - | 53 (53%) | 37 (28.2%) |
| Total tissue cores studied | 100 | 131 |

**Supplemental Table S2.** ALPPL2 expression in normal tissues: IHC study on a FFPE normal human tissue array. N: No staining.

| **Normal tissue type** | **Case 1** | **Case 2** |
|---|---|---|
| Adrenal gland | N | N |
| Bladder | N | N |
| Bone marrow | N | N |
| Eye | N | N |
| Breast | N | N |
| Cerebellum | N | N |
| Cerebral cortex | N | N |
| Fallopian tube | N | N |
| GI-Esophagus | N | N |
| GI-Stomach | N | N |
| GI-Small intestine | N | N |
| GI-Colon | N | N |
| GI-Rectum | N | N |
| Heart | N | N |
| Kidney | N | N |
| Liver | N | N |
| Lung | N | N |
| Ovary | N | N |
| Pancreas | N | N |
| Parathyroid | N | N |
| Pituitary gland | N | N |
| Placenta | Positive | Positive |
| Prostate | N | N |
| Skin | N | N |
| Spinal cord | N | N |
| Spleen | N | N |
| Striated muscle | N | N |
| Testis | N | N |
| Thymus | N | N |
| Thyroid | N | N |
| Tonsil | N | N |
| Uterus-cervix | N | N |
| Uterus-endometrium | N | N |

**Supplemental Table S3**. Affinity assessment of anti-ALPPL2 antibodies. Binding of yeast displaying scFvs to recombinant human ALPPL2 was determined by flow cytometry, and the apparent $K_D$ (M) was determined by curve-fitting of MFI values. The original M25 scFv, when displayed on yeast, did not show saturation binding over the range of concentrations tested (up to 40 nM), and $K_D$ was not determined. The two affinity improved clones (M25ADLF and M25FYIA) showed 0.582 nM and 0.446 nM apparent binding affinity to ALPPL2, respectively. Specificity of affinity-improved scFv was maintained, as evidenced by the lack of binding to human ALPI. No binding is indicated by "-".

| ScFv | Binding to ALPPL2 | Binding to ALPI |
|---|---|---|
| M25ADLF | $5.82 \times 10^{-10}$ | - |
| M25FYIA | $4.46 \times 10^{-10}$ | - |

**Supplemental Table S4**. Summary of IHC staining patterns of M25FYIA IgG1 on a frozen normal human tissue array. N: No staining.

| Tissue | Case 1 | Case 2 | Case 3 |
|---|---|---|---|
| Lymph node | N | N | N |
| Skeletal muscle | N | N | N |
| Prostate | N | N | N |
| Kidney | N | N | N |
| Liver | N | N | N |
| Lung | N | N | N |
| Stomach | N | N | N |
| Esophagus | N | N | N |
| Heart | N | N | N |
| Colon | N | N | N |
| Small intestine | N | N | N |
| Peripheral nerve | N | N | N |
| Smooth muscle | N | N | N |
| Cerebellum | N | N | N |
| Cerebrum | N | N | N |
| Ovary | N | N | N |
| Pancreas | N | N | N |
| Salivary gland | N | N | N |
| Adrenal gland | N | N | N |
| Placenta | Positive | Positive | Positive |
| Skin | N | N | N |
| Spinal cord | N | N | N |
| Spleen | N | N | N |
| Testis | N | N | N |
| Thymus | N | N | N |
| Thyroid gland | N | N | N |
| Ureter | N | N | N |
| Cervix | N | N | N |

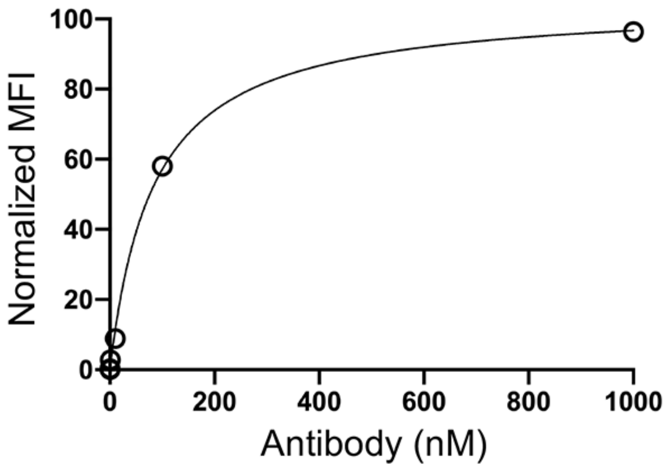


**Figure S1.** Apparent binding affinity of M25 scFv to cells expressing ALPPL2. Recombinant M25 scFv with a hexahistidine tag was produced from bacterial periplasmic space, affinity purified, biotin-labeled and incubated over a range of concentrations with HEK293 cells transfected with the ALPPL2 expression plasmid. Binding was measured by flow cytometry, and the apparent $K_D$ (~ 83 nM) was determined by curve-fitting MFI values.

**ALPI/ALPP_1**

MQGPWVLLLLGLRLQLSLGVIPAEEENPAFWNRQAAEALDAAKKLQPIQKVAKNLIIF
LGDGMGVSTVTAARILKGQKKDKLGPEIPLAMDRFPYVALSKTYNVDKHVPDSGAT
ATAYLCGVKGNFQTIGLSAAARFNQCNTTRGNEVISVMNRAKKAGKSVGVVTTTRV
QHASPAGTYAHTVNRNWYSDADVPASARQEGCQDIATQLISNMDIDVILGGGRKYM
FRMGTPDPEYPDDYSQGGTRLDGKNLVQEWLAKRQGARYVWNRTELMQASLDP
SVTHLMGLFEPGDMKYEIHRDSTLDPSLMEMTEAALRLLSRNPRGFFLFVEGGRID
HGHHESRAYRALTETIMFDDAIERAGQLTSEEDTLSLVTADHSHVFSFGGYPLRGSS
IFGLAPGKARDRKAYTVLLYGNGPGYVLKDGARPDVTESESGSPEYRQQSAVPLDE
ETHAGEDVAVFARGPQAHLVHGVQEQTFIAHVMAFAACLEPYTACDLAPPAGTTDA
AHPGRSVVPALLPLLAGTLLLLETATAP

**ALPI/ALPP_2**

MQGPWVLLLLGLRLQLSLGVIPAEEENPAFWNRQAAEALDAAKKLQPIQKVAKNLIL
FLGDGLGVPTVTATRILKGQKNGKLGPETPLAMDRFPYLALSKTYNVDRQVPDSAA
TATAYLCGVKANFQTIGLSAAARFNQCNTTRGNEVISVMNRAKQAGKSVGVVTTTR
VQHASPAGTYAHTVNRNWYSDADMPASARQEGCQDIATQLISNMDIDVILGGGRKY
MFPMGTPDPEYPADASQNGIRLDGKNLVQEWLAKRQGARYVWNRTELMQASLDP
SVTHLMGLFEPGDMKYEIHRDSTLDPSLMEMTEAALRLLSRNPRGFFLFVEGGRID
HGHHESRAYRALTETIMFDDAIERAGQLTSEEDTLSLVTADHSHVFSFGGYPLRGSS
IFGLAPGKARDRKAYTVLLYGNGPGYVLKDGARPDVTESESGSPEYRQQSAVPLDE
ETHAGEDVAVFARGPQAHLVHGVQEQTFIAHVMAFAACLEPYTACDLAPPAGTTDA
AHPGRSVVPALLPLLAGTLLLLETATAP

**Figure S2.** Amino acid sequences of the two ALPI/ALPP fusion constructs. The leader sequence of ALPL (amino acids 1-19) are used in both constructs.  Grey: ALPI sequence including the leader. Rose: ALPP sequence including the leader.

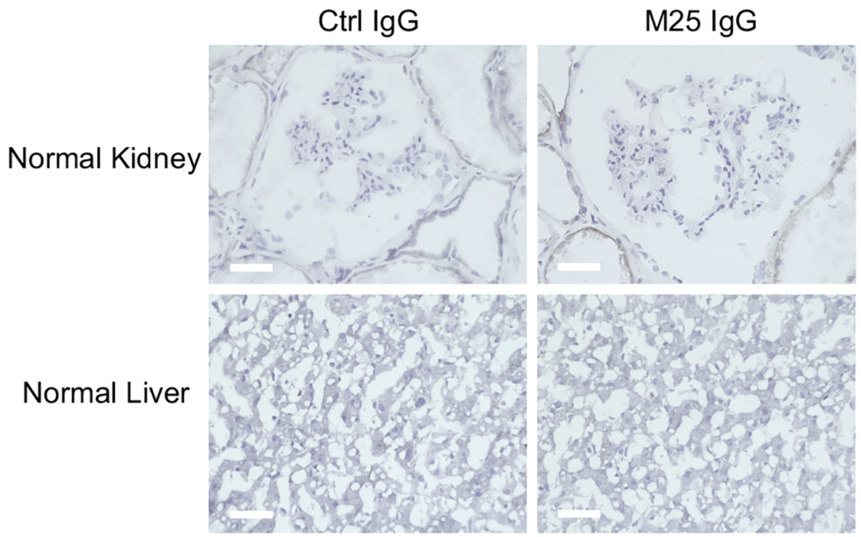


**Figure S3.** Additional M25 IgG1 IHC study of human frozen normal tissues. Besides tissue arrays, single sections from major organs such as kidney and liver were further stained to account for variables during tissue processing. Compared to control non-binding IgG, M25 IgG showed no positive staining on human frozen normal kidney and liver. Scale bar: 50 μm.

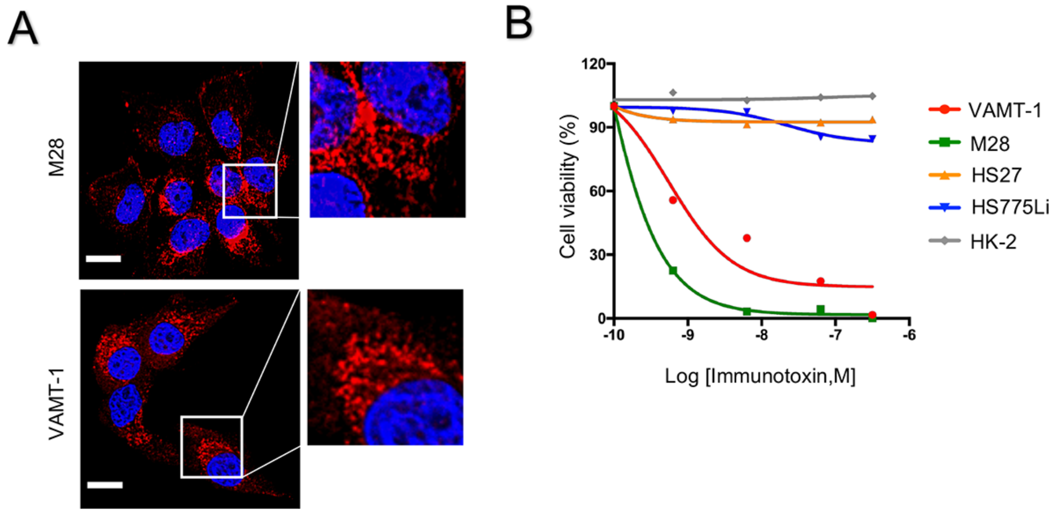


**Figure S4.** Tumor cell-specific internalization and cytotoxicity. (A) Internalization of M25 IgG1 into mesothelioma lines M28 and VAMT-1. Immunofluorescence microscopy study was performed after 24h of incubation. Scale bar: 30 μm. (B) The M25 IgG1-saporin immunotoxin killed both mesothelioma cell lines (M28 and VAMT-1), but not normal cell lines (HS27, HK-2, and HS775Li).

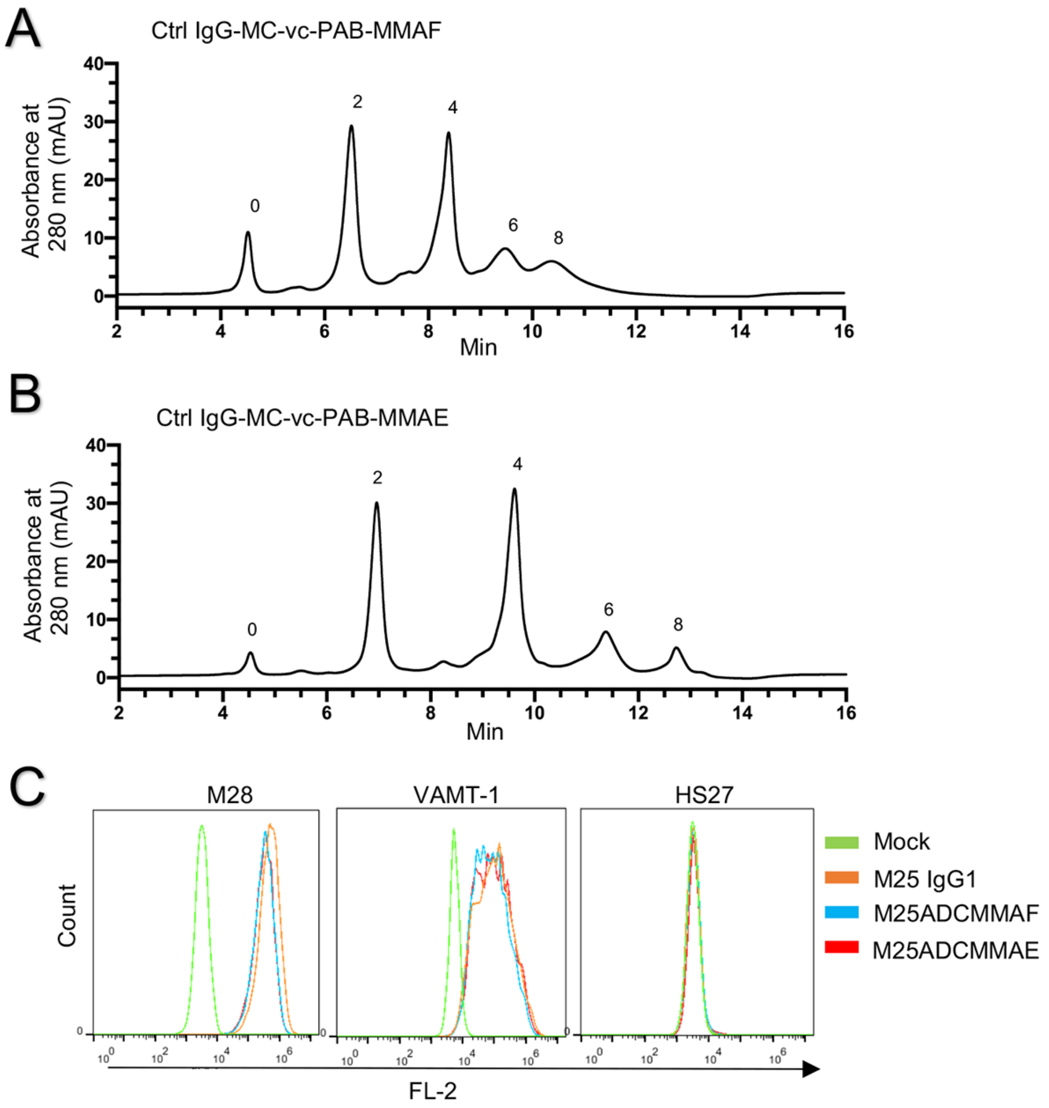


**Figure S5.** Characterization of anti-ALPPL2 ADC. (A) HIC analysis of MMAF-conjugated Ctrl IgG1 (Ctrl IgG-MC-vc-PAB-MMAF or CtrlADCMMAF). (B) HIC analysis of MMAE-conjugated Ctrl IgG1 (Ctrl IgG-MC-vc-PAB-MMAE or CtrlADCMMAE) (C) M25 ADC cell binding assessment. M25 IgG1 and M25 ADCs showed equivalent binding to mesothelioma cells, suggesting minimal impact of drug conjugation on antibody-antigen interaction.

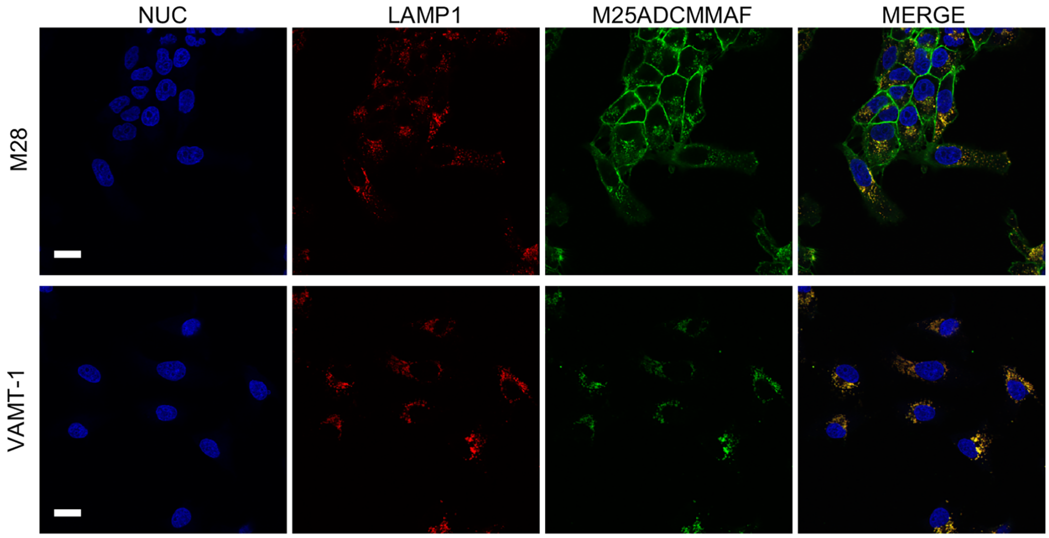


**Figure S6.** Subcellular localization of anti-ALPPL2 ADC in mesothelioma cells. Confocal microscopy study was performed after 24h of incubation to determine M25ADCMMAF co-localization with lysosome marker LAMP1. Upper panel: M28; Lower panel: VAMT-1. Nuc: nucleus. Scale bar: 30 μm.

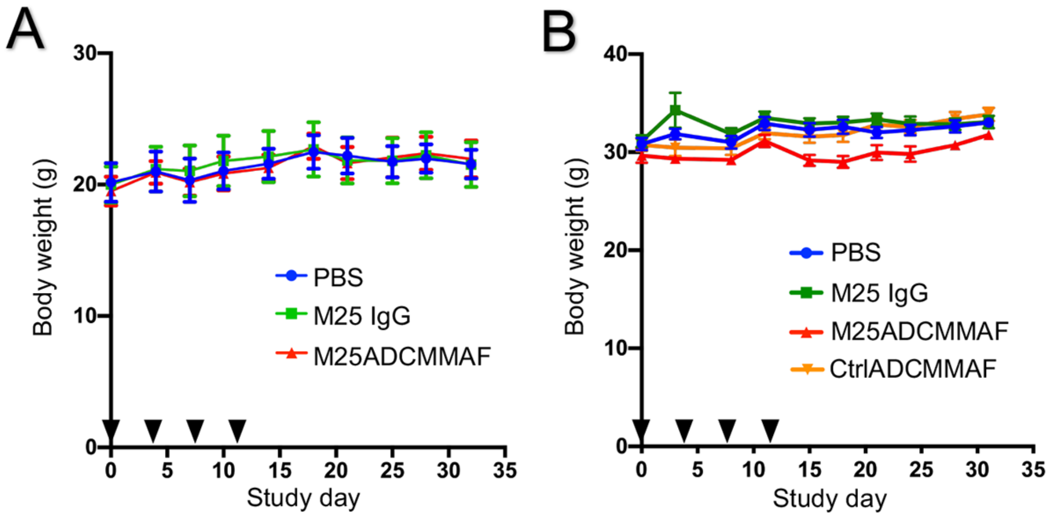


**Figure S7.** Body weight assessment. (A) Body weights of Nude mice bearing M28 xenografts (~250 $mm^3$) treated with M25ADCMMAF, CtrlADCMMAF as described in Figure 5B. (B) Body weights of NSG mice bearing M28 xenograft (~150 $mm^3$) treated with M25ADCMMAF, CtrlADCMMAF, vehicle only (PBS) and naked antibody (M25 IgG1) as described in Figure 5C. Injection days are indicated by black triangles.

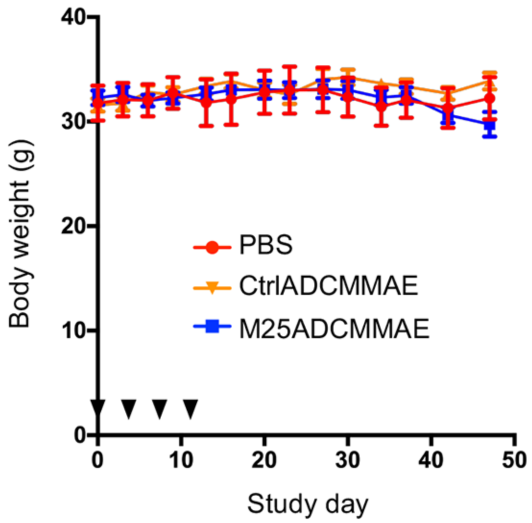


**Figure S8** Body weight assessment. Body weights of NSG mice bearing large-sized (~500 $mm^3$) M28 xenografts treated with M25ADCMMAE, CtrlADCMMAE and vehicle only (PBS) as described in Figure 5D. Injection days are indicated by black triangles.

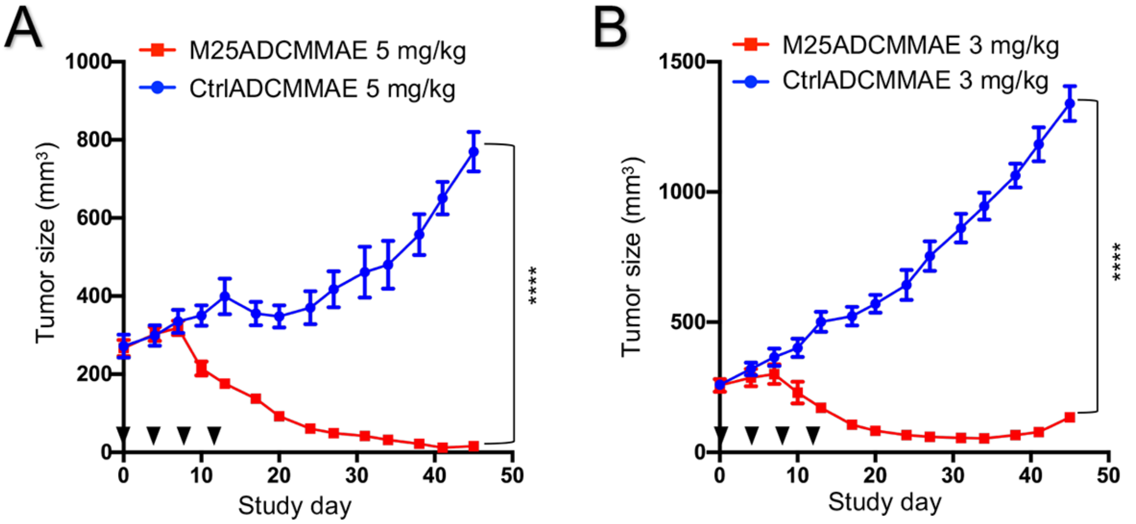


**Figure S9.** Statistical analysis of tumor volumes. (A) NSG mice bearing M28 xenografts (~250 mm$^3$) were treated with M25ADCMMAE or CtrlADCMMAE at 5 mg/kg every four days for total of 4 doses. Injection days are indicated by black triangles. Quantification of tumor sizes shows significant difference (**** $p < 0.0001$) at the end of the experiment. Student's t test, unpaired two-tailed. (B) NSG mice bearing M28 xenograft (~250 mm$^3$) were treated with M25ADCMMAE or CtrlADCMMAE at 3 mg/kg every four days for total of 4 doses. Injection days are indicated by black triangles. Quantification of tumor sizes shows significant difference (**** $p < 0.0001$) at the end of the experiment. Student T test, unpaired two-tailed.

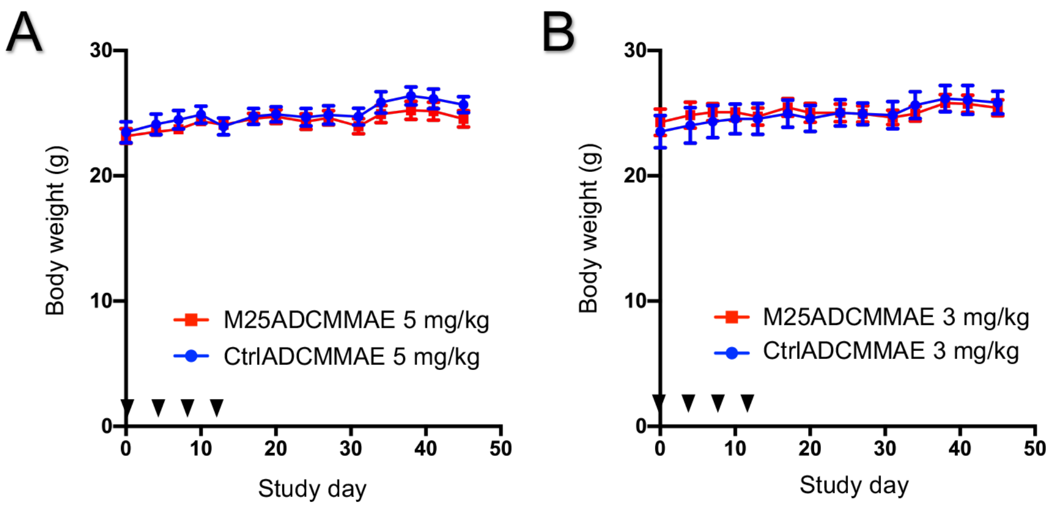


**Figure S10.** Body weight assessment. (A) Body weights of NSG mice treated with M25ADCMMAE or CtrlADCMMAE at 5 mg/kg. (B) Body weights of NSG mice treated with M25ADCMMAE or CtrlADCMMAE at 3 mg/kg. Injection days are indicated by black triangles.

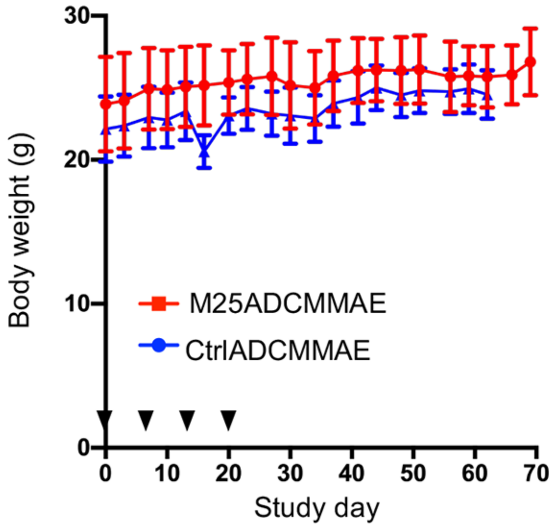


**Figure S11.** Body weight assessment of dually grafted mice. Injection days are indicated by black triangles.
